# 5-axis CNC micro-milling machine for three-dimensional microfluidics

**DOI:** 10.1101/2024.06.07.597629

**Authors:** Mitchell J. C. Modarelli, Devin M. Kot-Thompson, Kazunori Hoshino

## Abstract

The gold standard of microfluidic fabrication techniques, SU-8 patterning, requires photolithography equipment and facilities and is not suitable for 3D microfluidics. A 3D printer is more convenient and may achieve high resolutions comparable to conventional photolithography, but only with select materials. Alternatively, 5-axis CNC micro-milling machines can efficiently prototype structures with high resolutions, high aspect ratios, and non-planar geometries from a variety of materials. These machines, however, have not been catered for laboratory-based, small-batch microfluidics development and are largely inaccessible to researchers. In this paper, we present a new 5-axis CNC micro-milling machine specifically designed for prototyping 3D microfluidic channels, made affordable for research and laboratories. The machine is assembled from commercially available products and custom-build parts, occupying 0.72 cubic meters, and operating entirely from computer aided design (CAD) and manufacturing (CAM) software. The 5-axis CNC micro-milling machine achieves sub-µm bidirectional repeatability (≤0.23 µm), machinable features <20 µm, and a work volume of 50 x 50 x 68 mm. The tool compatibility and milling parameters were designed to enable fabrication of virtually any mill-able material including metals like aluminum, brass, stainless steel, and titanium alloys. To demonstrate milling high resolution and high aspect ratios, we milled a thin wall from 360 brass with a width of 18.1 µm and an aspect ratio of ∼50:1. We also demonstrated fabricating molds from 360 brass with non-planar geometries to create PDMS microfluidic channels. These included a channel on a 90° edge and a channel on a rounded edge with a 250-µm radius of curvature. Our 5-axis CNC micro-milling machine offers the most versatility in prototyping microfluidics by enabling high resolutions, geometric complexity, a large work volume, and broad material compatibility, all within a user-friendly benchtop system.

## 1.0 Introduction

Microfluidics are a crucial tool across multiple disciplines of science and research.^1,2^ As efforts in the field of microfluidics increase, techniques and equipment have become more accessible to researchers, driving new ideas, methods, and applications of microfluidics in academia and industry.^3^

Many microfluidic fabrication techniques exist because each offers its own advantages and disadvantages^4^. Choosing the best fabrication technique depends on the desired resolution, geometry, material, cost, production time, and application of the microfluidic device in development.^4^ For example, SU-8 photolithography or silicon micropatterning have produced the highest achievable resolutions, creating sub-µm structures.^4,5^ However, the necessary equipment, clean room, trained operator, and production time make these techniques costly for prototyping, customization, or small batch production of microfluidic devices.^4,6^ SU-8 photolithography or silicon micropatterning are also limited to select materials and planar geometries.^4,7^ As a counter example, 3D printing offers very fast and low-cost production of microfluidics with complex 2D and 3D geometries.^4,8,9^ The minimal equipment, working environment, and computer aided design required for 3D printing make this technique accessible and affordable.^8^ However, achieving high precision and surface quality, especially on slanted geometries, remains challenging for 3D printing, and the material choice is limited by each 3D printing method.^4,8,10,11^

Collectively, current fabrication techniques for microfluidic devices provide a broad range of solutions. However, there is a growing demand in the biomedical field for microfluidics using biocompatible materials and 3D shapes that current techniques do not meet.^12,13^ We propose using 5-axis CNC micro-milling as a fabrication method for microfluidic devices with material diversity and 3D shapes. Currently, 3-axis milling has been reported for prototyping and fabricating microfluidics, but this technique is limited in geometric complexity.^14-18^ 5-axis systems expand the capabilities of traditional 3-axis systems by including two rotational axes, allowing the workpiece to be milled from virtually any position. 5-axis systems can mill overhanging geometries, curved surfaces, and inclined features. These systems can perform continuous milling operations to avoid manually rotating and realigning the workpiece. Continuous operations provide high material removal rates by keeping the tool in constant contact with the workpiece.^19,20^ This also improves the accuracy and surface quality of parts since the cutting tool can move along 5 axes. 5-axis CNC micro-milling systems can produce uniquely curved channels, amorphous surfaces, angled through holes, and non-planar chambers.^21,22^ Therefore, these systems have potential applications in microfluidics. However, 5-axis CNC micro-milling machines are largely reserved for high-end industries.^23,24^ These machines are not catered to the fabrication of microfluidics and are not readily accessible to researchers.^25^ Therefore, we have designed a unique 5-axis CNC micro-milling machine for affordable prototyping, customization, and small batch production of microfluidic devices. Our machine prioritizes four critical factors: (1) resolution, (2) geometric complexity, (3) material compatibility, and (4) costs of the equipment, facilities, and fabrication.

Our first design objective was to achieve high resolution structures within tens of µm in size and aspect ratios comparable to the best techniques. For example, chlorine ICP etching is capable of aspect ratios >32:1, demonstrated by Volatier, et al.^26^ UV laser cutting has a resolution in the range of 30 μm,^27^ and 39 ± 15-µm paper based microchannels have been demonstrated using CO2 laser cutting.^28^ In this paper, we demonstrate a 5-axis CNC micro-milling machine that competes with these resolutions. Our machine also includes a system of three microscopic cameras for real time control of milling operations. The machining surface and tool condition can be monitored in-situ, providing real-time feedback to optimize machining parameters.

Our second objective was a high design flexibility to create geometrically complex features. 3D printing has set a high standard for microfluidics with geometric complexity, such as 3D printed microfluidic mixers, channels on curved surfaces, and integrated electronic sensors demonstrated by Su, et al.^9^ Unlike any other microfluidic fabrication technique, 5-axis CNC micro-milling can compete with 3D printing’s design flexibility because of its degrees of freedom, and with 3D printing’s easy operation because of its similar implementation of computer aided design and manufacturing. Our 5-axis CNC micro-milling machine has five axes with sub-µm precision and uses CAD and CAM software to produce complex 3D microfluidic channels.

Our third objective was for our machine to be compatible with the most versatile selection of materials. The operating parameters of CNC milling are highly customizable, such as the spindle speed, feed rate, tool diameter, tool length, and radial and axial depths of cut. This allows nearly any metal, polymer, composite, ceramic, or other hard material to be milled as a microfluidic channel or a master for a microfluidic mold.^20,29^ Stainless steel, for example, is biocompatible and important in biomedical applications but cannot be machined with traditional MEMS techniques. Unlike MEMS processes, 5-axis CNC micro-milling does not involve mask fabrication or lithography, broadening usable materials and providing faster turnaround time necessary for prototyping microfluidics.^20,30^

Our fourth objective was for our 5-axis CNC micro-milling to not require expensive facilities such as a clean room or specialized equipment. In this paper, we demonstrate a benchtop 5-axis CNC micro-milling machine made entirely from commercially available products and custom-made parts, occupying 0.72 cubic meters. We have also reduced the costs of fabrication to predictable tool wear, raw materials, and power consumption.

We have designed a new 5-axis CNC micro-milling machine catered to the fabrication of microfluidic devices that achieves (1) high resolution, (2) complex 3D geometries, (3) versatile material compatibility, and (4) efficient fabrication costs. By prioritizing these four critical factors, we are making 5-axis CNC micro-milling machines affordable to researchers and applicable to microfluidics research.

## 2.0 Experimental

### 2.1 Design of the 5-axis CNC micro-milling machine

The 5-axis CNC micro-milling machine is constructed from commercially available products, including the computer modelling and operating software, firmware, milling station hardware, and system of microscopic cameras for in-situ monitoring. The system loop diagram is shown in Figure 1, and the data flow diagram is shown in Figure 2.

**Fig. 1.**
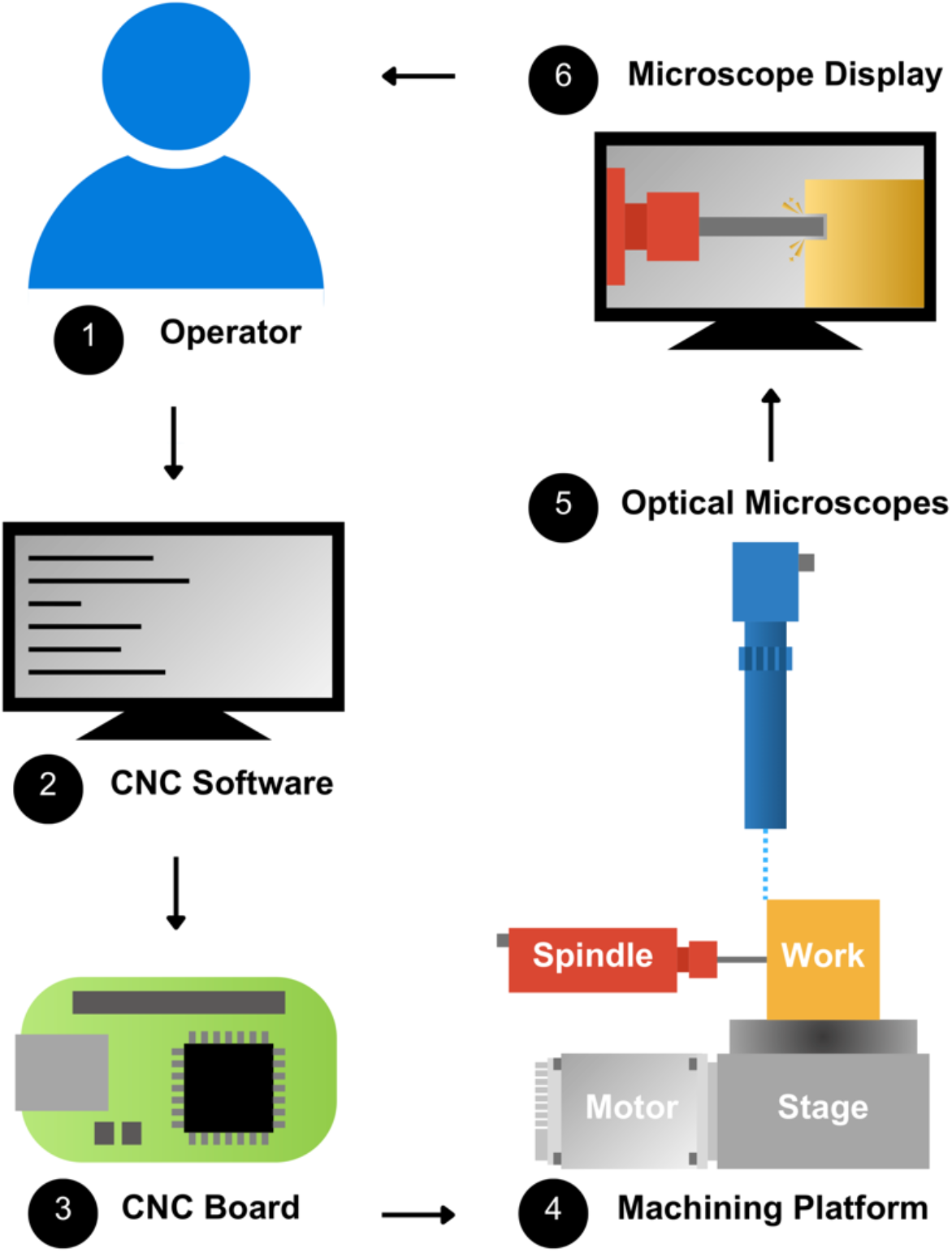
System loop diagram: (1) The operator provides G-code instructions through (2) the CNC software. (3) The CNC board receives the G-code instructions and controls (4) the servomotors driving the linear and rotational stages to move the spindle and workpiece during milling operations. (5) The optical microscopes provide (6) in-situ feedback to the operator, and (1) the operator provides new instructions.

**Fig. 2.**
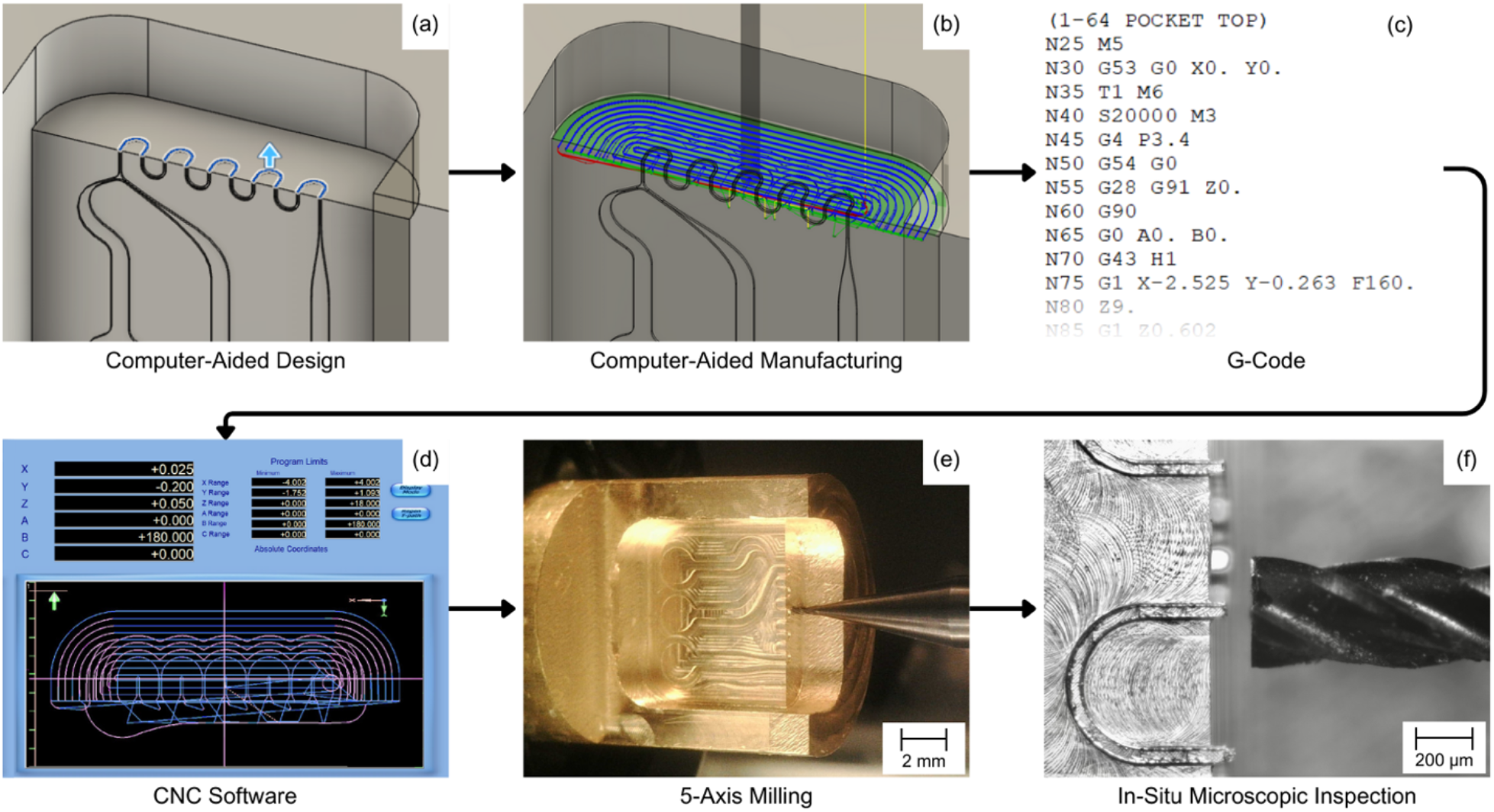
Data flow diagram: (a) The 3D model is created using Fusion 360 CAD; (b) then the manufacturing toolpath is designed with Fusion 360 CAM and (c) exported as G-code; (d) the G-code is imported into Mach3 on a desktop computer, which is used to (e) operate the 5-axis mill; (F) a system of three optical microscopes image the milling process and provide real-time feedback on the desktop computer.

#### 2.1.1 Spindle

The mill spindle (Sfida MS01, Minitor, Tokyo, Japan) is a brushless motor spindle with ceramic angular bearings and a stainless-steel body. It is designed for small-diameter end mills (30 µm – 4 mm), achieving a 1-μm accuracy and speeds between 5,000 and 60,000 rpm. This range of rotational speeds allows for micro-milling resins, plastics, woods, and a wide variety of metals, including aluminium, brass, stainless steel, and titanium alloys. The spindle is driven by an electric motor and uses compressed-air cooling to prevent heating in the motor and keep the bearings clean.

#### 2.1.2 Air supply

We supply the spindle’s cooling mechanism using an air compressor (8010SPC, California Air Tools, Inc., San Diego, CA) and a pressure regulator (Sfida MT01CP, Minitor, Tokyo, Japan). The compressor supplies a typical pressure of 0.42 MPa to the regulator. Then, the regulator reduces the pressure to a constant 0.25 MPa, specified by the spindle manufacturer, and supplies air to the spindle during operation.

#### 2.1.3 Workpiece lubrication and cleaning

The air pressure regulator also supplies an air nozzle above the spindle. The air nozzle has an adjustable air pressure gauge for various working conditions. When working with metals, the air nozzle deposits lubricating oil onto the workpiece to reduce friction and chatter while evacuating milled chips. Then, a vacuum fixed below the endmill collects milled chips during fabrication. The air nozzle and vacuum are fixed to the spindle stage to avoid collisions with the other stages while allowing their full range of motion.

#### 2.1.4 Stages and servomotors

The 5-axis CNC micro-milling machine is equipped with three linear axes (X, Y, Z) and two rotational axes (A, B) creating a work volume of 50 x 50 x 68 mm (X, Y, Z). The X– and Z-axes use a high precision dual-axis ball-screw linear stage (XYRB-4040, Dover Motion, Boxborough, MA) with ±25 mm ranges of motion. The Y-axis uses a high precision ball-screw linear stage (LMB-200, Dover Motion) with a ±34 mm range of motion and an increased axial load capacity to carry the A– and B-axis stages (Figure 3). The A– and B-axes use rotary stages with crossed roller bearing and a 45:1 worm/gear ratio (RTR-4, Dover Motion). The A-axis has a ±90° range of motion, and the B-axis range is unlimited. All five axes are controlled by step-and-direction servomotors (ClearPath SDSK Series, CPM-SDSK-2311P-EQN, Teknic, Inc., Victor, NY) with resolutions of 6,400 steps/rev. The servomotors have embedded drivers and are powered by a low voltage (75 VDC) power supply (IPC-5, Teknic, Inc).

**Fig. 3.**
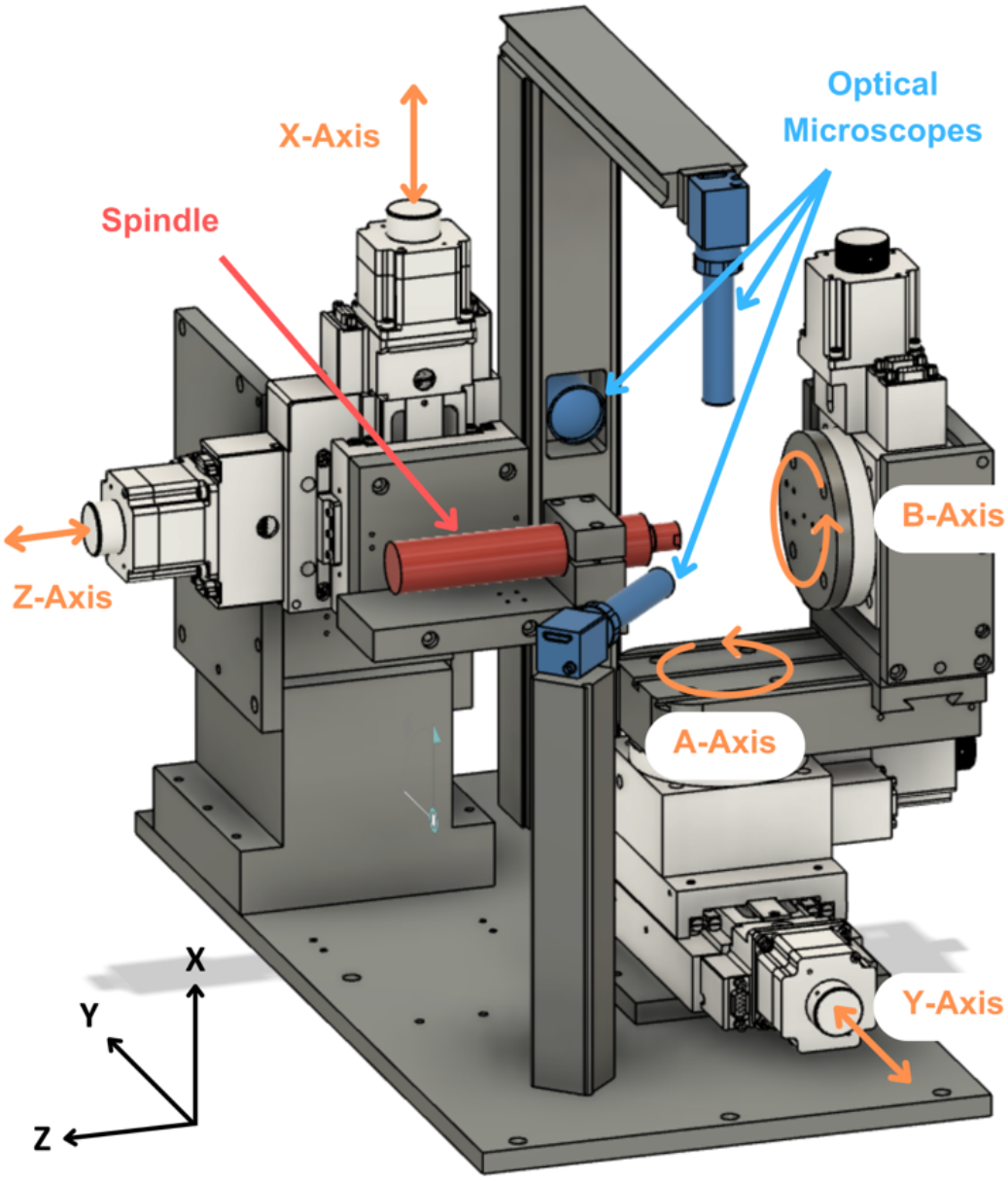
Illustration of 5-axis CNC micro-milling station, including the spindle (red), linear and rotational stages (white), and microscopes (blue).

#### 2.1.5 CNC controller

We used the Pokeys57CNC to control the step-and-direction signal-driven servomotors. The Pokeys57CNC can control up to 8 motors, which is sufficient for our 5-axis system, and it is compatible with Mach3 CNC software. Mach3 operates on a typical computer as a 5-axis CNC machine controller with a customizable interface and G-code display, functioning primarily by communicating with the firmware controller and processing G-code. G-code was either written manually or generated using Fusion 360, a manufacturing software.

#### 2.1.6 Microscopic optical inspection

Three microscopes (Edmund Optics, Barrington, NJ) observe the workspace for in-situ, multidirectional, microscopic inspection during milling and for tool and part alignments. The microscopes along the X-axis (model 67317) and arbitrary-axis (model 33110) can be adjusted in X-, Y-, and Z-directions. The arbitrary-axis microscope has an additional roll adjustment, and the microscope along the Y-axis (model 67316) is fixed. All three microscopes are displayed on the PC monitor for real-time optical feedback. The arrangement of the spindle, stages, and optical microscopes of the 5-axis CNC micro-milling machine is illustrated in Figure 3.

#### 2.1.7 Axis alignment

The three microscopes were used to optically align the tool to the absolute origin of the machine. When working with 3-axis machines, the X-, Y-, and Z-origins can be set arbitrarily by zeroing the machine’s axes. However, with the inclusion of two rotational axes in our 5-axis setup, there exists an absolute origin where all five axes intersect. To ensure the µm-level precision of the machine during multi-axis operations, we aligned the endmill to the absolute origin using multidirectional microscopic inspection.

We first milled a thin brass pillar and fixed this pillar to the B-axis. The A-axis stage was rotated +90° so that the pillar was aligned with the Y-axis. The Y-axis camera (Model 33110) was used to observe the position of the brass pillar on the X-Z plane. By rotating the B-axis 180° we found the center of rotation, which determined our X– and Z-origins. The endmill was aligned to the X– and Z-origins and zeroed.

The brass pillar was removed and fixed to the A-axis stage so that the pillar was aligned with the vertical X-axis. The X-axis camera (Model 67317) was used to observe the position of the brass pillar on the Y-Z plane. By rotating the A-axis 180° we found the center of rotation, which determined our Y-origin and confirmed our Z-origin. The endmill was aligned to the Y-origin and zeroed.

When exchanging endmills, only the Z-axis required realignment. The current endmill was placed at the origin, and using one microscope, the position of the origin was recorded on the computer monitor. The Z-axis was retracted and the endmill was exchanged. Then the endmill was aligned with the origin recorded on the monitor, and the Z-axis was zeroed.

### 2.2 Fabrication of high aspect ratio walls

#### 2.2.1 Milling of thin walls

The 5-axis CNC micro-milling machine we present is capable of features with high aspect ratios. To demonstrate this, we milled thin walls from 360 brass. We started by preparing a stock using a 1/16”-diameter, 3-flute, solid carbide, square endmill operating at a 12,000-rpm spindle speed, 90 mm/min feed rate, and 0.050 mm depth of cut. The result was a rectangular wall 1 x 1.5 x 4 mm (W x H x L).

To create thin walls, we milled two adjacent trenches. We used a 100 µm-diameter, 2-flute, solid carbide, square endmill with a 1 mm neck length and 300 µm length of cut. We milled at a 30,000-rpm spindle speed, 0.5 mm/min feed rate, and 150 µm depth of cut. After the first trench was milled, the brass stock was cleaned with air, then wetted with acetone. Once the acetone evaporated, a drop of cyanoacrylate glue (commonly known as super glue) was applied to the trench to secure the thin wall. The cure time for the super glue was 24 minutes. Then a second trench was milled with the same parameters, creating the thin wall between trenches.

Once the thin walls were complete, the brass stock was removed and cleaned with compressed air, then submerged in acetone for 10 minutes to remove lubricating oil and glue. We did not use sonication because the thin wall was delicate. Lastly, the thin wall was cleaned in a plasma cleaner (PDC-32G, Harrick Plasma) for 60 seconds. Figure 4 shows SEM (ZEISS Crossbeam 340) images of the thin wall. The width of the wall varied slightly; we determined the maximum width to be 18.1 µm. The wall height was 900.4 providing an aspect ratio of 49.7:1.

**Fig. 4.**
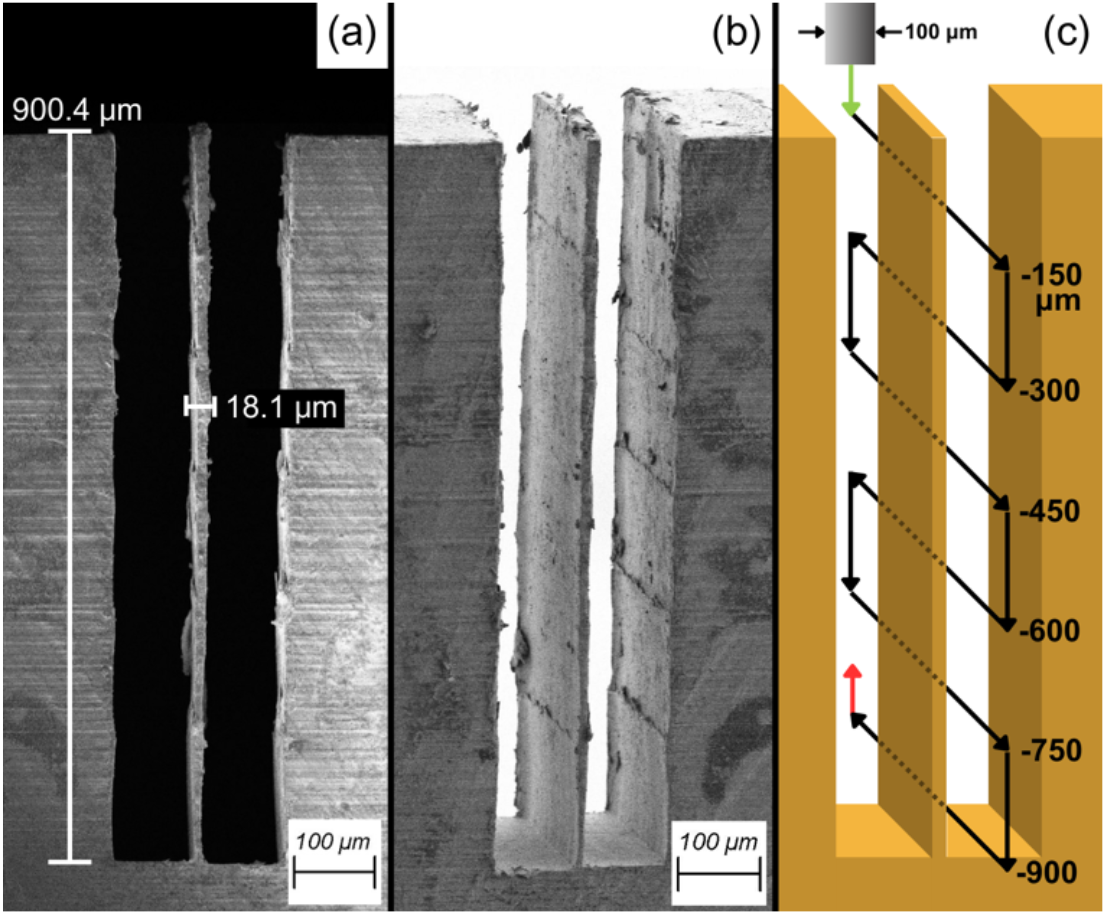
SEM images of thin wall with 49.7:1 aspect ratio: (a) side view at 180x magnification; (b) angled view at 175x magnification; (c) illustration of toolpath.

#### 2.2.2 Multi-axis milling of thin walls

To demonstrate multi-axis milling we also created a series of thin walls with side-milling features. Figure 5 illustrates the milling process to create three thin walls with a perpendicular slot and a slot at a 60° angle to the walls. To create the side-milled walls (Fig. 6) we started by milling the perpendicular channel from the brass stock, then milling the channel at a 60° angle. We recentered the stock and milled the three thin walls using the process detailed in Section 2.2.1.

### 2.3 Fabrication of brass molds

#### 2.3.1 3D modelling

We used Fusion 360 to create 3D models of microfluidic channel master molds and generate their manufacturing toolpaths. Three molds, namely planar, 90° edge, and 250-µm radius channels, illustrated in Figure 7, were created to demonstrate channels requiring multi-axis machining. The three channels were designed with two inputs and one output and a square cross section of 50 x 50 µm. Channel 1 is a planar channel with a winding pattern of nine 180° bends with a 250-µm radius (Fig. 7a). Channel 2 is the same channel with an additional 90° edge incorporated along the direction of the winding channel (Fig. 7b). Channel 3 is the same channel as Channel 1 with an additional rounded edge (250-µm radius) incorporated along the direction of the winding channel (Fig. 7c). All three channels have the same total length of 9 mm.

**Fig. 5.**
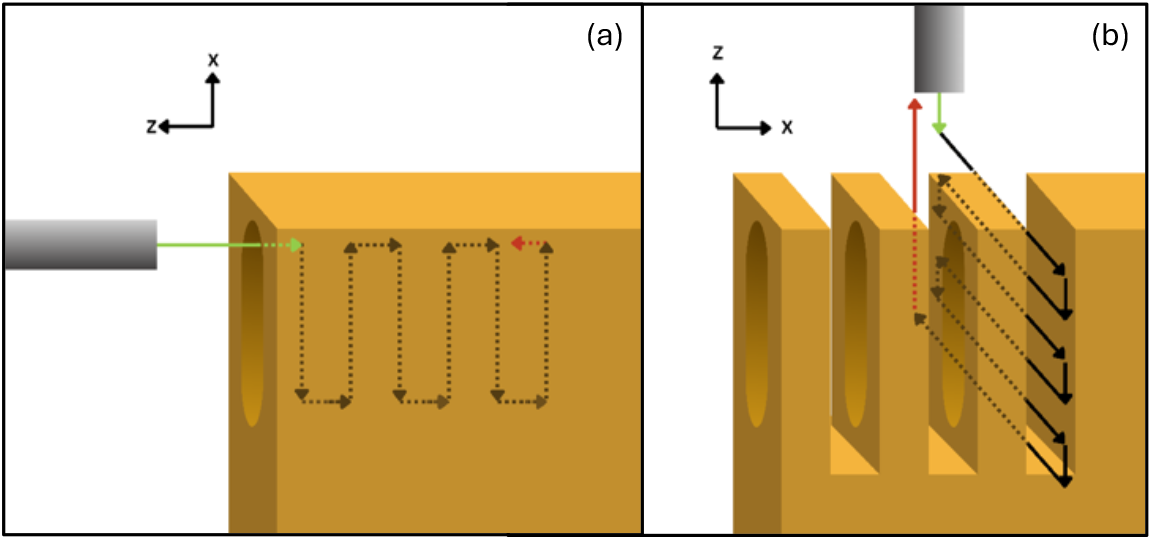
Illustration of the milling toolpath for the thin walls with multi-axis milling features; (a) milling perpendicular slot; (b) milling thin walls.

**Fig. 6.**
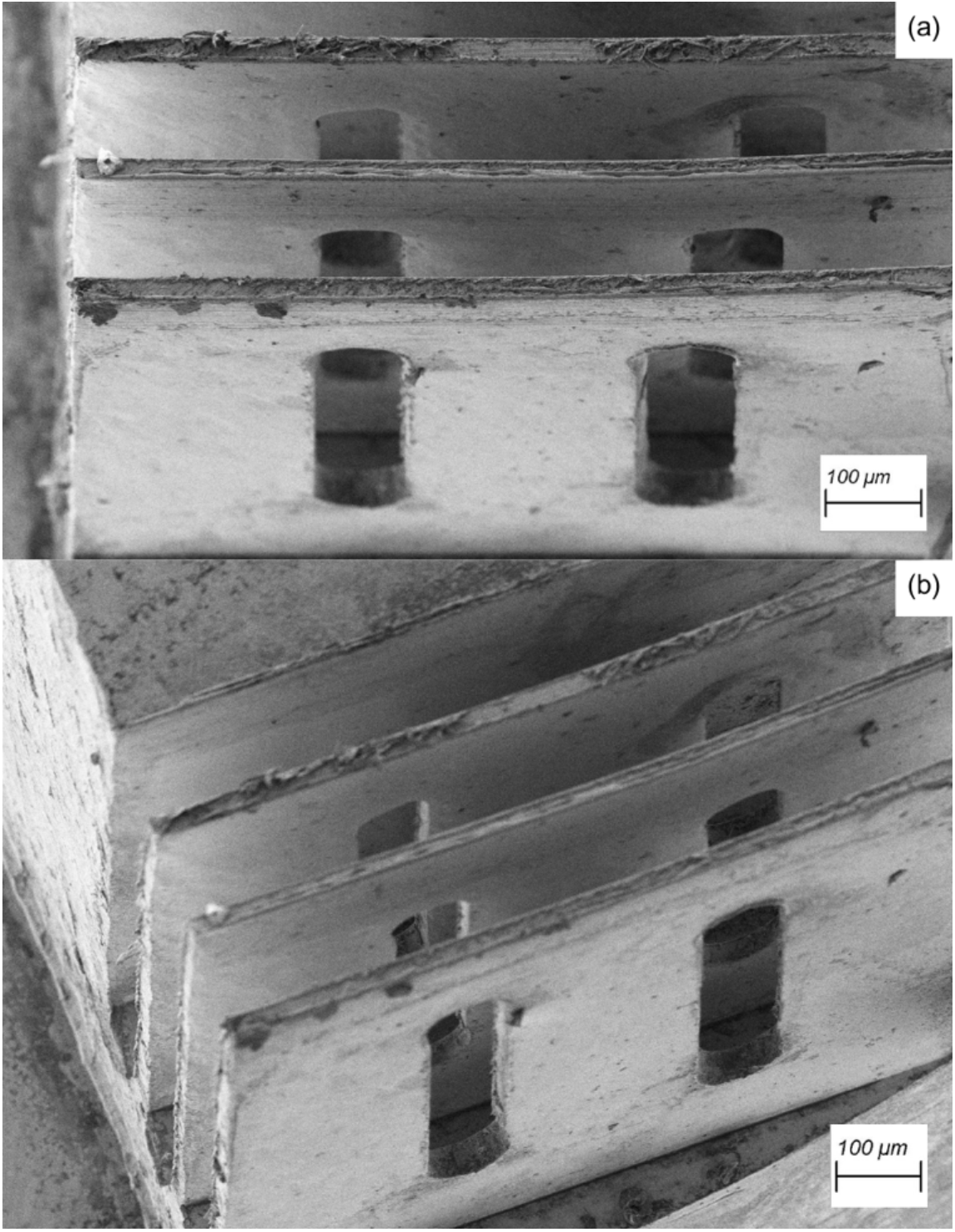
SEM images of thin walls with side slots: (a) 250x magnification; (b) 200x magnification.

**Fig. 7.**
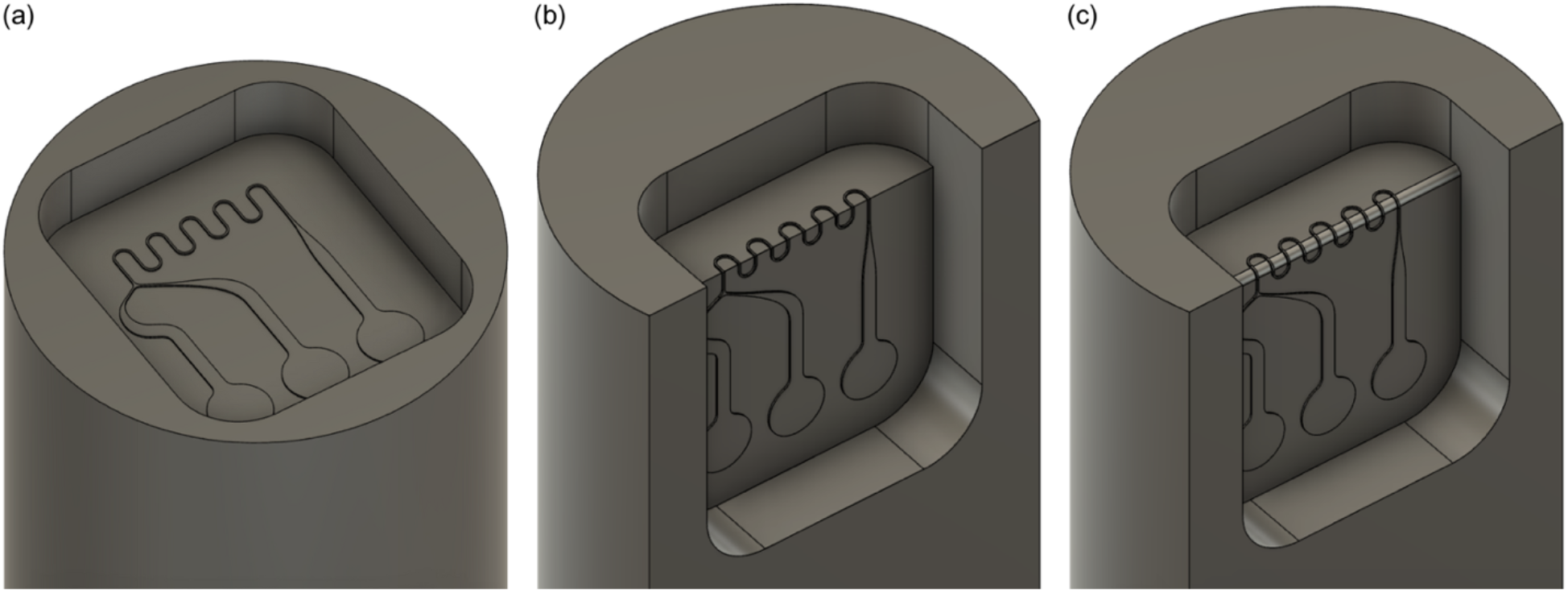
CAD models of (a) Channel 1 with a planar pattern, (b) Channel 2 with a 90° edge, and (c) Channel 3 with a rounded edge with a 250-µm radius of curvature.

#### 2.3.2 Manufacturing toolpaths

Toolpaths for the three microchannel molds were designed using Fusion 360. Milling the molds required roughing and finishing processes. In the roughing process we utilized a 1/8”-diameter, 3-flute, solid carbide, square endmill operating at a 12,000-rpm spindle speed, 180 mm/min feed rate, and 0.100 mm depth of cut. The roughing process involved an initial surfacing of the stock top followed by the removal of a 1.93-mm deep pocket, leaving 70 µm of depth for the finishing process. Channels 2 and 3 both required a 90° y-axis rotation to mill both faces of the microfluidic channels during the roughing process.

In the finishing process, we utilized a 1/64”-diameter, 3 flute, solid carbide, square endmill operating at a 20,000-rpm spindle speed, 45 mm/min feed rate, and 0.001 mm depth of cut. The finishing process involved a 20-µm deep surfacing within the mold pocket, followed by the cleanout of the 50-µm deep channels. Channels 2 and 3 both required a 90° y-axis rotation to mill both faces of the microfluidic channels. Channel 3 also required incremental rotation around the y-axis during the finishing process to create the curved channel walls.

Once the roughing and finishing toolpaths were generated, the milling processes were simulated in Fusion 360 to ensure the machine avoided collisions during and between operations. Then we output G-code compatible with Mach3, our CNC software.

#### 2.3.3 Milling

The three microfluidic channel molds were made from alloy 360 brass rods with 1/2”-diameter. The 5-axis CNC milling machine was aligned with the roughing endmill and the brass stock was fixed to the B-axis rotational stage, oriented along the Z-axis. Lubricating oil was used during operation to reduce friction and chatter between the tool and stock while assisting the removal of milled chips. After milling, the brass molds were cleaned with compressed air, then submerged in acetone for 10 minutes during sonication. Lastly, the molds were cleaned in a plasma cleaner (PDC-32G, Harrick Plasma) for 60 seconds.

#### 2.3.4 PDMS molding

We created polydimethylsiloxane (PDMS) microfluidic channels from the brass master molds using Sylgard 184 Silicone Elastomer Kit at a 10:1 ratio of base to curing agent. Once mixed, the uncured PDMS was degassed in a vacuum for 30 minutes. Then the PDMS was poured into the brass master molds and degassed for an additional 30 minutes. These PDMS molds were the bottom components of the microfluidic channels. Top components were made from a rectangular acrylic master mold. In this mold, one edge was given 250-µm fillet to match the curvature of Channel 3. The molds were covered and placed in an oven at 70°C for 3 hours.

After removing the molds from the oven and allowing to cool to room temperature, the cured PDMS molds were bonded using plasma treatment. Both top and bottom components were placed in the plasma cleaner for 45 seconds. The surfaces of the top and bottom components were wetted with isopropanol, immediately placed together, and placed in the oven at 70°C for 60 minutes. The PDMS channels were removed from the oven and allowed to cool to room temperature. The channels were then placed in a larger acrylic mold and submerged in uncured PDMS to prevent separation of the top and bottom PDMS components and to allow for better handling of the channels. After curing in the oven under the same conditions, The PDMS channels were removed and cooled. Holes with 1.5 mm diameters were punched into the PDMS for setup of the fluid tubes. Fabrication of the PDMS microchannels is illustrated in Figure 8.

**Fig. 8.**
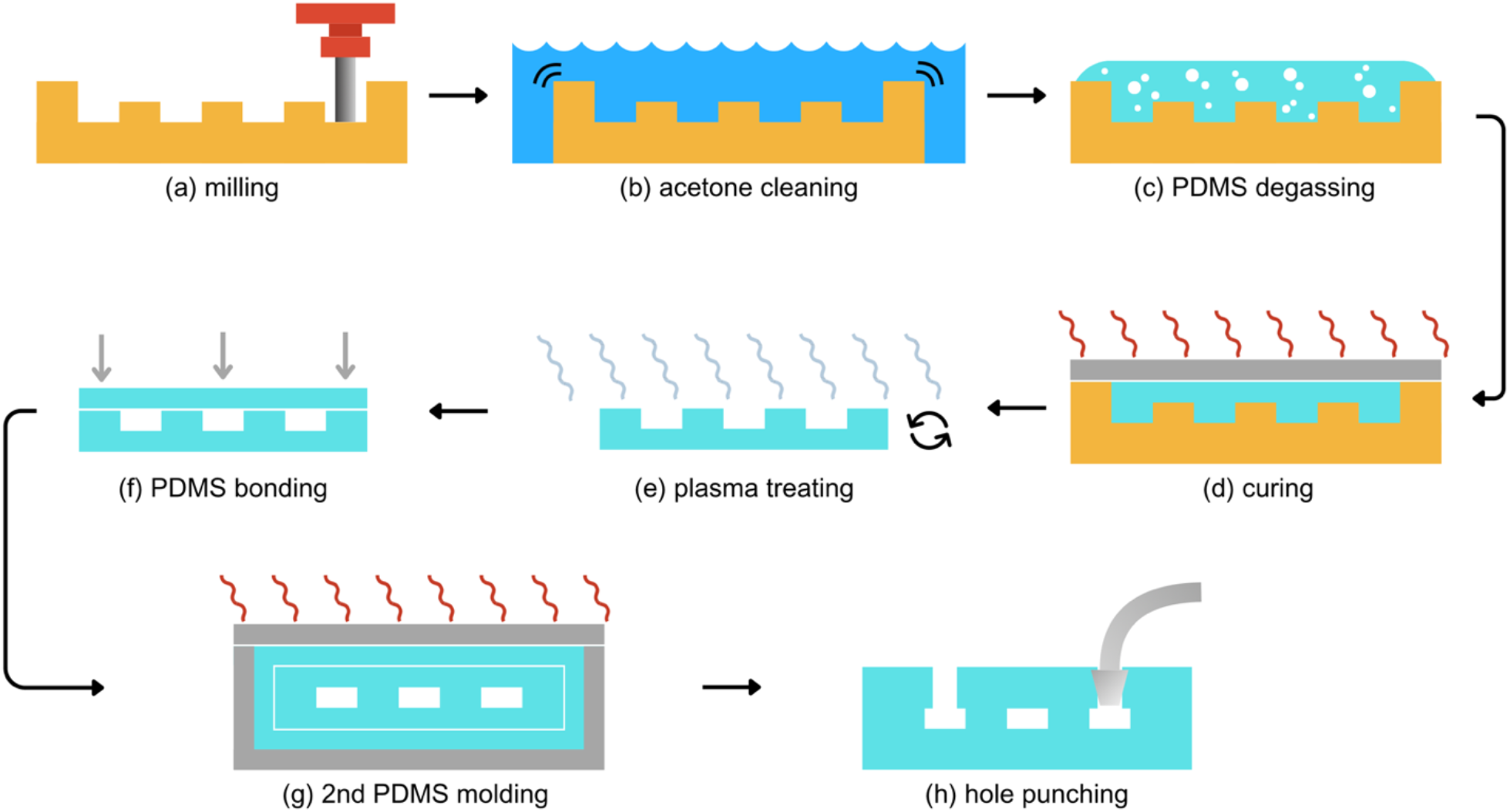
Illustrated fabrication of PDMS microfluidic channels; (a) mill brass molds; (b) clean with sonication in acetone; (c) fill molds with uncured PDMS and degas; (d) cover and cure PDMS at 70°C for 3 hours; (e) plasma treat cured PDMS; (f) bond PDMS microchannel; (g) embed microchannel in large PDMS mold; (h) hole punch cured microchannel for tube assembly.

## 3.0 Results and discussion

### 3.2 Bidirectional repeatability of the milling machine

The bidirectional repeatability of the 5-axis CNC micro-milling machine is a combined result of the stages, servomotors, and assembly. To test the bidirectional repeatability of the five axes, each axis was measured independently for 20 iterations of movement along its maximum range of motion, limiting the A-axis to ±90° of rotation and the B-axis to ±360° of rotation. A needle was fixed to each axis, and the needle’s position during each iteration of movement was photographed with a microscope (33110, Edmund Optics, Barrington, NJ) and measured using ImageJ.

Absolute deviations of the linear and rotational axes are shown in Figures 9 and 10, respectively. The standard deviations for the 5 axes are listed as the bidirectional repeatability in Table 1. Over 20 iterations of cyclic movement, the X-, Y-, and Z-axes had positional ranges (max–min) less than 1 µm, and the A– and B-axes had angular ranges less than 0.005°. The linear axes also demonstrated sub-µm bidirectional repeatability. The Y-axis showed the highest standard deviation, likely because of its higher range of motion and the higher weight of the assembly. The A– and B-axes both showed standard deviations of 0.001° (Table 1).

**Table 1.** Calculated bidirectional repeatability of five axes.

| Axis | Bidirectional Repeatability |
| --- | --- |
| X | 0.12 $\mu\text{m}$ |
| Y | 0.23 $\mu\text{m}$ |
| Z | 0.16 $\mu\text{m}$ |
| A | 0.001° |
| B | 0.001° |

**Fig. 9.**
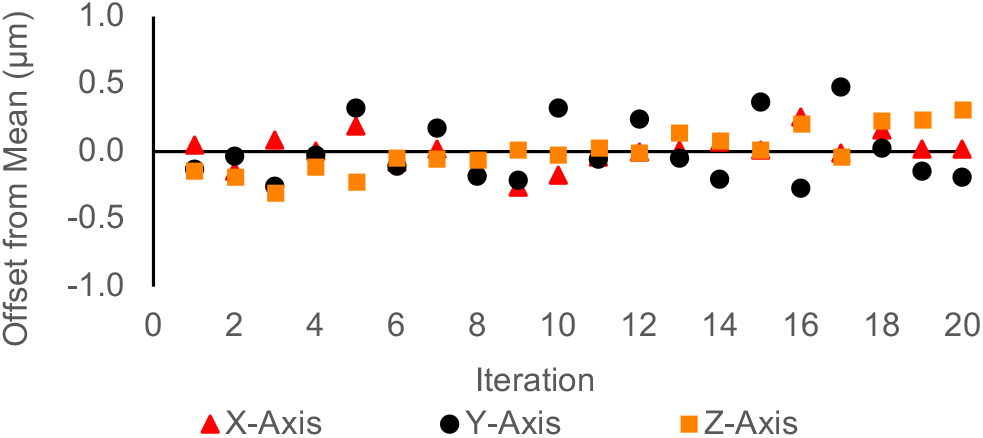
Measured absolute deviation of linear axes.

**Fig. 10.**
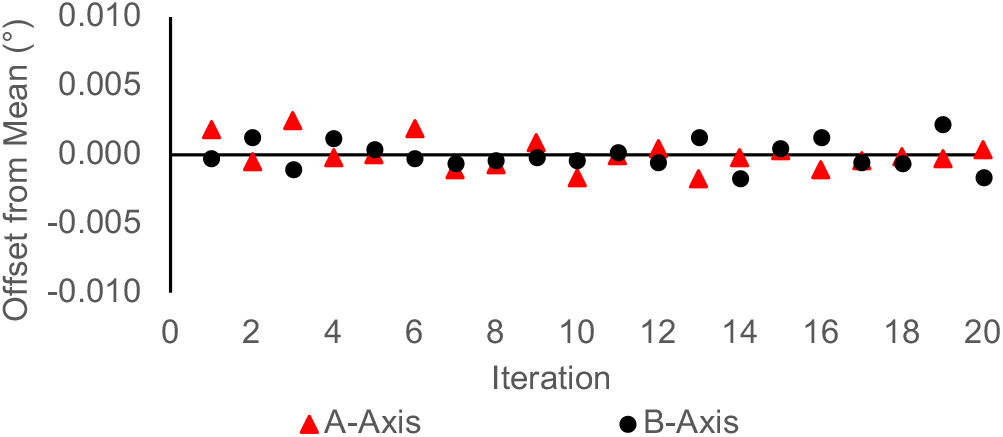
Measured absolute deviation of rotational axes.

### 3.2 Dimensional analysis of microfluidic channel molds

We took scanning electron microscope images of the three brass master molds to determine the dimensional precision along the channel and dimensional accuracy compared to the computer aided design. Figure 11 shows SEM images of the three channel master molds, and Figure 12 charts the width measurements of the channels, each measured at 40 locations using ImageJ software. Table 2 provides the accuracy (averages) and precision (standard deviations) of the channel dimensions. The design width of the channels was 50 µm, and the widths of Channels 1, 2, and 3 averaged 53.9 µm, 55.1 µm, and 63.7 µm, respectively. Channel 1 may have the highest dimensional accuracy because it had the simplest toolpath, only milling on a single plane.

**Table 2.** Accuracy and precision of milled microfluidic channel master mold widths.

| Channel | 1 | 2 | 3 |
| --- | --- | --- | --- |
| Average ($\mu\text{m}$) | 53.9 | 55.1 | 63.7 |
| SD ($\mu\text{m}$) | -3.3 | 4.5 | -3.5 |

**Fig. 11.**
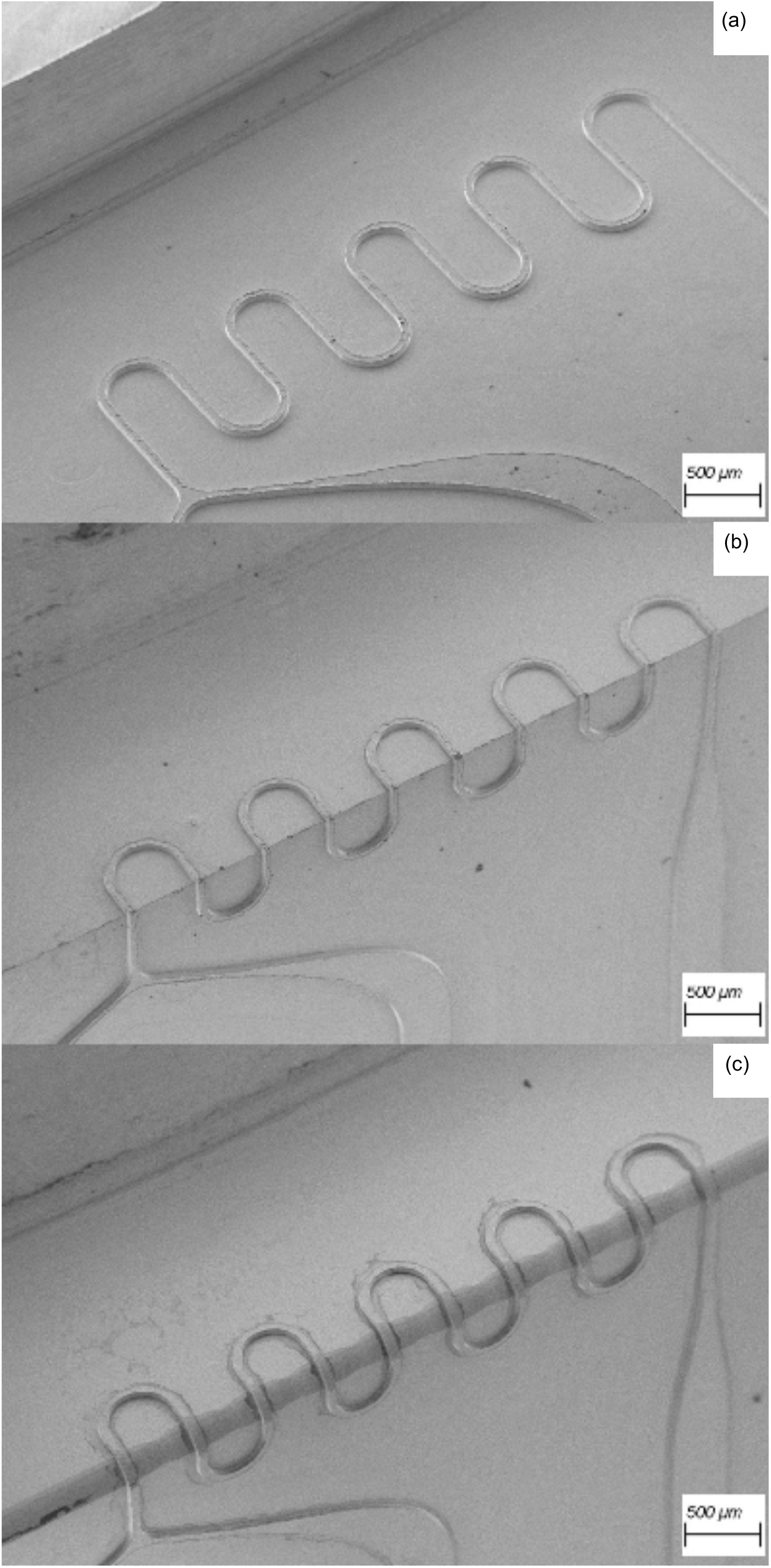
SEM images of brass master molds at 45x magnification: (a) Channel 1 (planar); (b) Channel 2 (90° edge); (c) Channel 3 (250-µm radius edge).

**Fig. 12.**
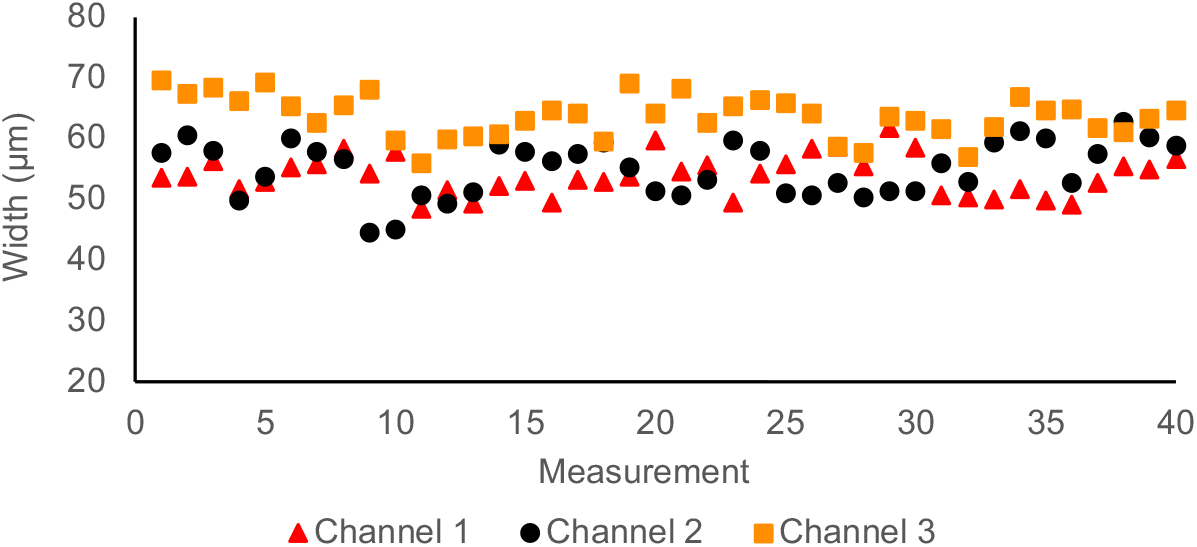
Width measurements of Channels 1, 2, and 3.

The dimensional error is likely a result of the milling process rather than the accuracy or repeatability of the machine. The main causes of geometric inaccuracies are tool deflection, tool runout, and machining chatter.^31,32^ These factors will vary the position of the endmill radially but not axially. Because of the low strength and stiffness of small endmills, such as the 1-64”-diameter tool used, material removal rates must be maintained by retaining high spindle speeds above 20,000 rpm and slow, smooth, and continuous feed rates.^31^ However, higher feed rates generate more chatter. Therefore, we can improve the dimensional accuracy and precision of the channels by optimizing the spindle speed and feed rate given our work material and tool diameter; and other milling parameters can be manipulated including the radial (Ae) and axial (Ap) depths of cut, endmill coating, helix angle, flute count, lubrication, etc. As discussed in the previous section, we achieved sub 20 µm, very high aspect ratio (∼50:1) structures. Further width reduction is easily achievable with a lower aspect ratio (∼1:1) channel as commonly used in microfluidics.

All three channels demonstrated average widths greater than the design. This variation can be corrected in the computer aided manufacturing by adding an additional finishing toolpath to refine the channel widths. While increasing production time, additional finishing passes can optimize the surface and dimensional accuracy of the channels. The SEM images also revealed burring along the edges of Channels 1, 2, and 3. Burring on our microfluidic channel is shown in Figure 13a. Burring is caused by plastic and elastic deformation, predominately a result of the work material properties and the endmill geometry, deflection, runout, and chatter.^31^ Because burrs deteriorate the precision, function, and performance of machined parts, removing and even preventing burrs has become a complicated and extensively researched subject.^31^ To deburr our channel, we added a contour finishing pass on the sides followed by a pass on the top surface using the 1/64”-diameter, 3 flute, solid carbide, square endmill. We used a spindle speed of 20,000-rpm and reduced the feed rate to 5 mm/min. Figure 13b shows the results of deburring.

**Fig. 13.**
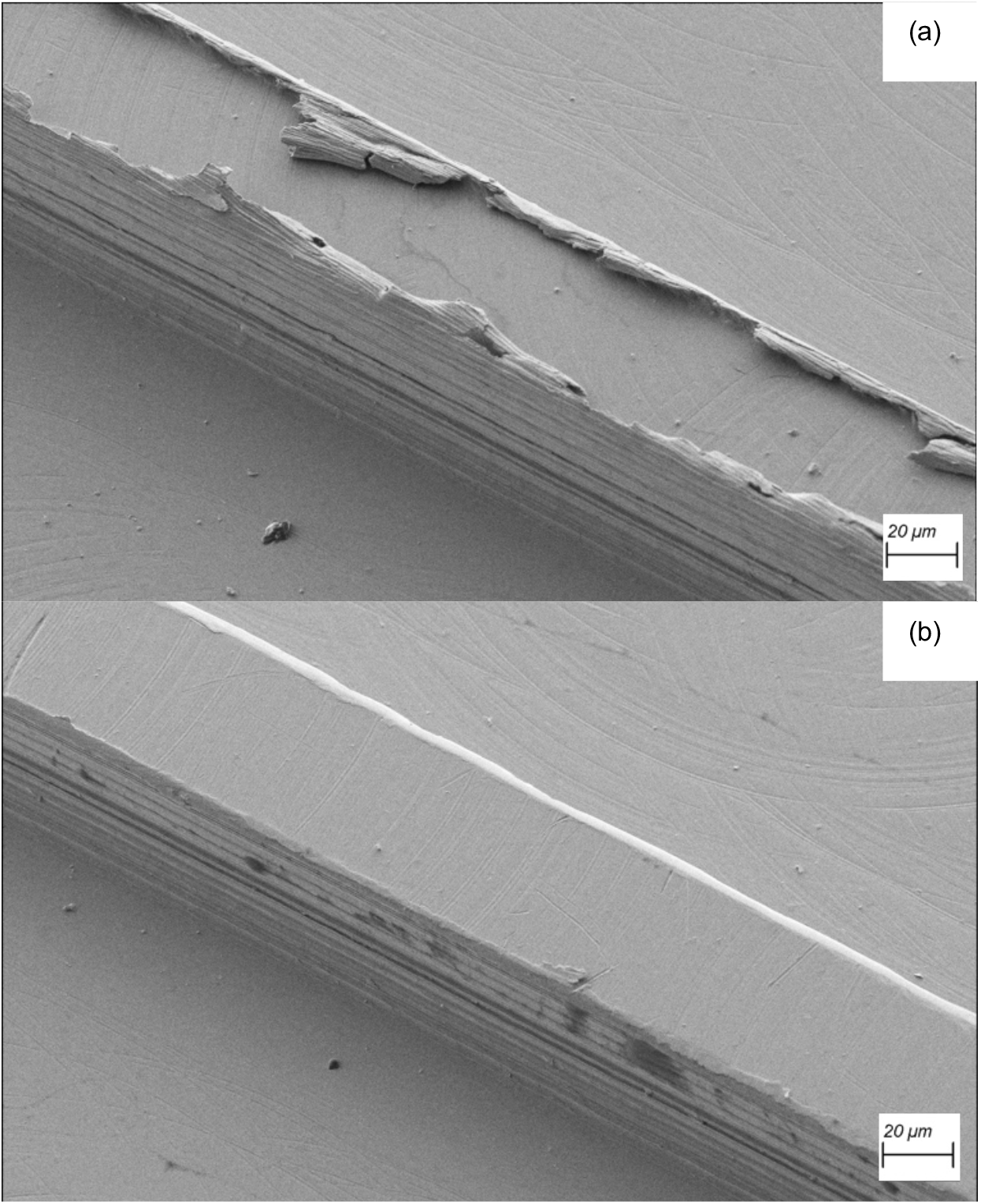
SEM image of 360 brass microfluidic channel at 1000x magnification (a) without and (b) with deburring.

### 3.3 Channel testing

The three PDMS microfluidic channels were qualitatively assessed by supplying the two inputs with deionized water containing red food dye. Figure 14 shows the function of all three channels. Figure 14b reflects the 90° edge of Channel 2, and Figure 14c shows the rounded edge of Channel 3 with a 250-µm radius of curvature.

**Fig. 14.**
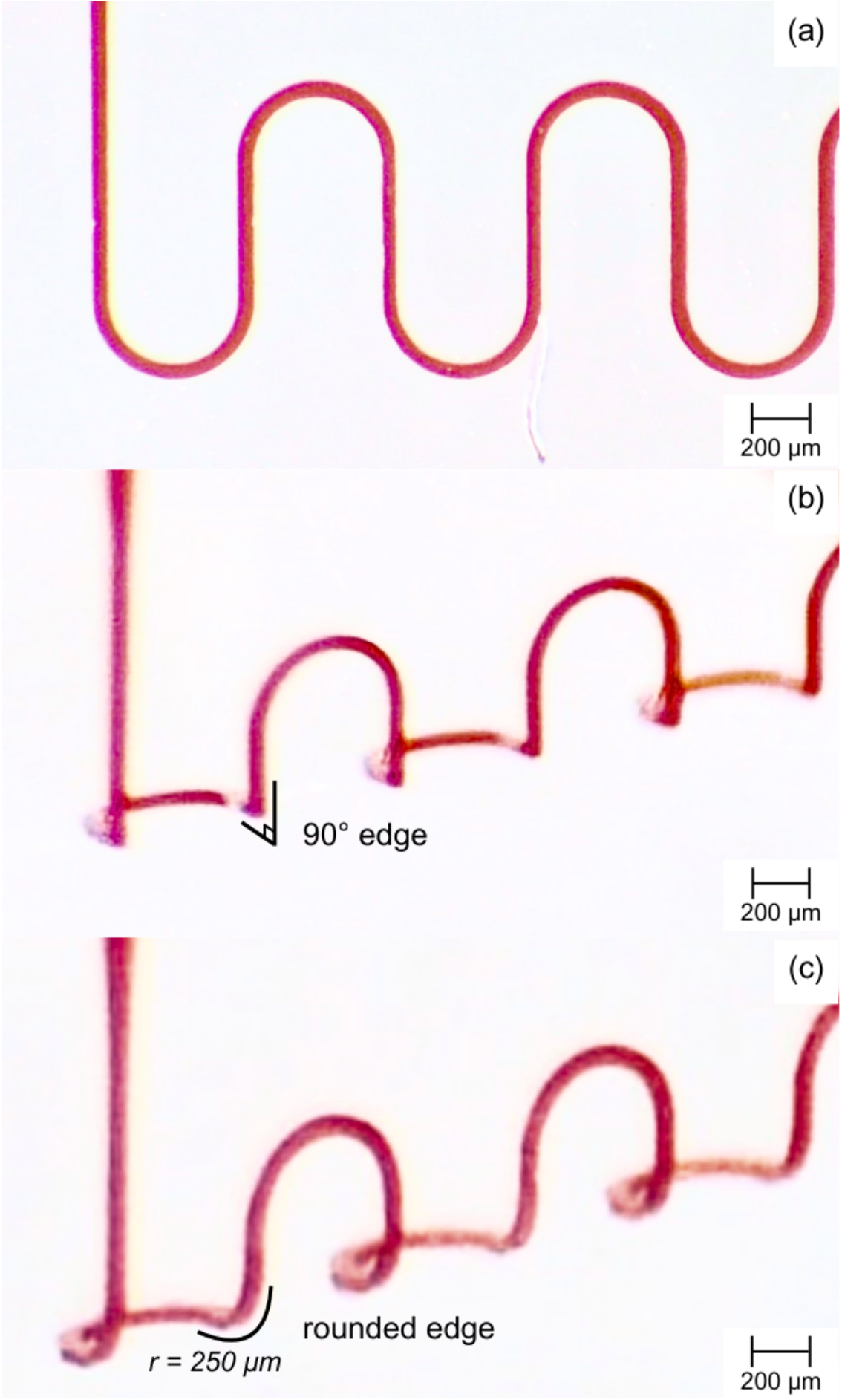
Optical images of PDMS microfluidic channels: (a) Channel 1 (planar); (b) Channel 2 (90° edge); (c) Channel 3 (250-µm radius edge).

## Conclusions

We designed, built, and tested a 5-axis CNC micro-milling machine catered to prototype microfluidic channels with 3D flow paths. The machine achieved (1) high resolution, (2) complex 3D geometries, (3) versatile material compatibility, and (4) efficient fabrication costs.

We demonstrated the high resolution of the 5-axis CNC micro-milling machine by milling a thin wall with a width <20 µm and an aspect ratio of 49.7:1. Then we demonstrated milling small features with geometries unusual to microfabrication. We milled a series of three thin walls with two slots cutting through the surfaces of all three walls. Additionally, we milled microfluidic channel (width ∼50 µm) molds with non-planar geometries. One channel demonstrated a 90° edge at the intersection of two perpendicular planes, and a second channel demonstrated a rounded edge with a 250-µm radius of curvature.

The 5-axis CNC micro-milling machine was designed with versatile material compatibility and low operating costs. The motor spindle has a 1-µm accuracy and speeds between 5,000 and 60,000 rpm, compatible with resins, plastics, woods, and metals. The 5-axis CNC micro-milling machine reduces fabrication costs for small-batch research and development by its design freedom and operating requirements. Because fabrication only requires CAD and CAM, operating the machine is simple and user-friendly. CAD and CAM also allow users the most design freedom to reduce production time and material costs under their unique milling criteria. The 5-axis CNC micro-milling machine also requires minimal equipment and working conditions. The entire system only occupies 0.72 cubic meters and does not require expensive facilities or special equipment.

Our 5-axis CNC micro-milling machine allows researchers to work autonomously to prototype complex 3D microfluidic channels and molds. We demonstrate sub-µm bidirectional repeatability of all five axes (≤0.23 µm) and a resolution below 20 µm. Making 5-axis CNC micro-milling accessible to research and laboratory environments will provide new opportunities to advance the development of microfluidic devices and benefit the multiple disciplines of science and research utilizing microfluidics.

## Author Contributions

Conceptualization: M.J.C.M., D.M.K.-T. and K.H.; Data Curation: M.J.C.M.; Funding Acquisition: K.H.; Investigation: M.J.C.M., D.M.K.-T. and K.H; Methodology: M.J.C.M, D.M.K.-T. and K.H.; Project Administration: K.H.; Resources: K.H.; Supervision: K.H.; Writing – original draft: M.J.C.M.; Writing – review & editing: M.J.C.M. and K.H.

## Acknowledgements

We thank the National Institute of Undersea Vehicle Technology (NIUVT) for funding support.

